# Temperature-dependent nanomechanics and topography of bacteriophage T7

**DOI:** 10.1101/326520

**Authors:** Zsuzsanna Vörös, Gabriella Csík, Levente Herényi, Miklós Kellermayer

## Abstract

Viruses are nanoscale infectious agents which may be inactivated by heat treatment. Although heat inactivation is thought to be caused by the release of genetic material from the capsid, the thermally-induced structural changes in viruses are little known. Here we measured the heat-induced changes in the properties of T7 bacteriophage particles exposed to two-stage (65 °C and 80 °C) thermal effect by using AFM-based nanomechanical and topographical measurements. We found that exposure to 65 °C caused the release of genomic DNA due to the loss of the capsid tail which leads to a destabilization of the T7 particles. Further heating to 80 °C surprisingly led to an increase in mechanical stability due to partial denaturation of the capsomeric proteins kept within the global capsid arrangement.

## Importance

Even though the loss of DNA, caused by heat treatment, destabilizes the T7 capsid, it is remarkably able to withstand high temperatures with a more-or-less intact global topographical structure. Thus, partial denaturation within the global structural constraints of the viral capsid may have a stabilizing effect. Understanding the structural design of viruses may help in constructing artificial nanocapsules for the packaging and delivery of materials under harsh environmental conditions.

## Introduction

Viruses are remarkable nanoscale machineries that harbor a piece of genetic material within a proteinaceous capsule and, as obligatory parasites, are capable of efficiently fooling the host organism into manufacturing the viral structural elements that spontaneously reproduce the virus particle by self-assembly. Because of their biological, medical and even economical importance, the properties of viruses have been investigated by a wide array of experimental approaches. It has long been known that most viruses can be thermally inactivated (1–4). It is hypothesized that thermal virus inactivation is caused by the release of the genetic material, but the exact nature of the thermally-driven structural transitions within the viruses are little known. Differential scanning calorimetry and cryo-electron microscopy experiments have revealed a reversible structural transition at 53 °C limited to the hexamers of the HK97 bacteriophage (5). Heating the HK97 phage further results in the release of genomic DNA by not precisely known mechanisms, and heating even further to 80 °C results in an irreversible transition of thermal melting (6). In the case of bacteriophage *λ* heat-induced transitions at 68 °C and 87 °C have been assigned to the escape of DNA and irreversible melting, respectively (7). The simultaneous observation of capsid- and DNA-related events, however, have not so far been possible at the level of the individual virus particles. Previously a distinct thermal melting of the bacteriophage T7 has been documented (8–11). T7 is a non-enveloped, short-tailed icosahedral *E.coli* phage that contains a 40 kbp genomic DNA (12). Thermal melting, measured by following OD_260_ as a function of temperature, involves two major transitions related to DNA. The first transition occurs between 50–60 °C and is thought to correspond to the release of DNA from the capsid. This transition is accompanied by a marked loss of infectivity (13). A second transition is detected in the sample at temperatures above 80 °C and is related to DNA denaturation. Although high-resolution structural information is available on the protein capsid of T7 (14, 15), because the thermal melting experiments measure the structural changes of DNA only, the thermally-induced transitions within the protein components of the capsid remain unclear.

In recent years atomic force microscopy (AFM)-based nanomechanical experiments emerged as a sensitive tool to explore the properties of viruses (16–25). It has been shown that nanomechanical parameters such as stiffness and capsid breaking force may reveal molecular mechanisms underlying capsid maturation and the packaging, storage and release of genetic material.

Here we employed AFM to explore the nanomechanical and topographical changes in T7 bacteriophages exposed to two-stage thermal treatment (65 °C, 80 °C). We show that distinct changes in the nanomechanical properties of T7 occur upon heat treatment. Topographical analysis revealed the structural alterations that underlie the nanomechanical changes: 65 °C treatment leads to the release of genomic DNA due to the loss of the tail complex, and further heating to 80 °C leads, on one hand, to the appearance of large globular particles that likely correspond to disassembled capsids and, on the other hand, to a partial structural stabilization of the remaining capsids due most likely to rearrangements *via* partial denaturation of the capsomeric gp10A proteins.

## Results and Discussion

### Nanomechanics of heat-treated T7 phages

In the present work individual, surface-adsorbed T7 phage particles exposed to different temperatures (room temperature, 65 °C, 80 °C) were manipulated with AFM to reveal their nanomechanical properties and the thermally-induced changes in these properties. **Fig. 1** shows our results obtained on phages at room temperature (RT). After landing the AFM tip on the capsid surface force increased linearly to about 8 nN where a sharp transition marked by a sudden drop of force and corresponding to capsid breakage occurred. Upon pressing the AFM tip further, force fluctuated below 2 nN, then it began to rise sharply upon approaching the substrate surface. The retraction force trace was essentially featureless; therefore, a large force hysteresis was present indicating that the mechanical manipulation resulted in an irreversible conformational change (breakage) of the capsid. Although in the majority of the capsids similar force traces were recorded (**Fig. 1.c**), in a fraction of them we obtained traces with a significantly different, but reproducible, appearance (**Fig. 1.d**). In these traces the initial linear regime ended at about 2 nN (we refer to these as putative empty capsids, see below). In T7 capsids treated at 65 °C (**Fig. 2.a-b**) the force traces were similar to those seen in **Fig. 1.d**: capsid breakage occurred at about 2 nN, then force fluctuated around 2 nN before increasing abruptly upon approaching the substrate surface. In T7 capsids treated at 80 °C (**Fig. 2.c-d**) the overall appearance of the force traces was similar to that seen for the 65 °C samples, but capsid breakage and the following force fluctuation occurred at greater force levels.

**Figure 1.**
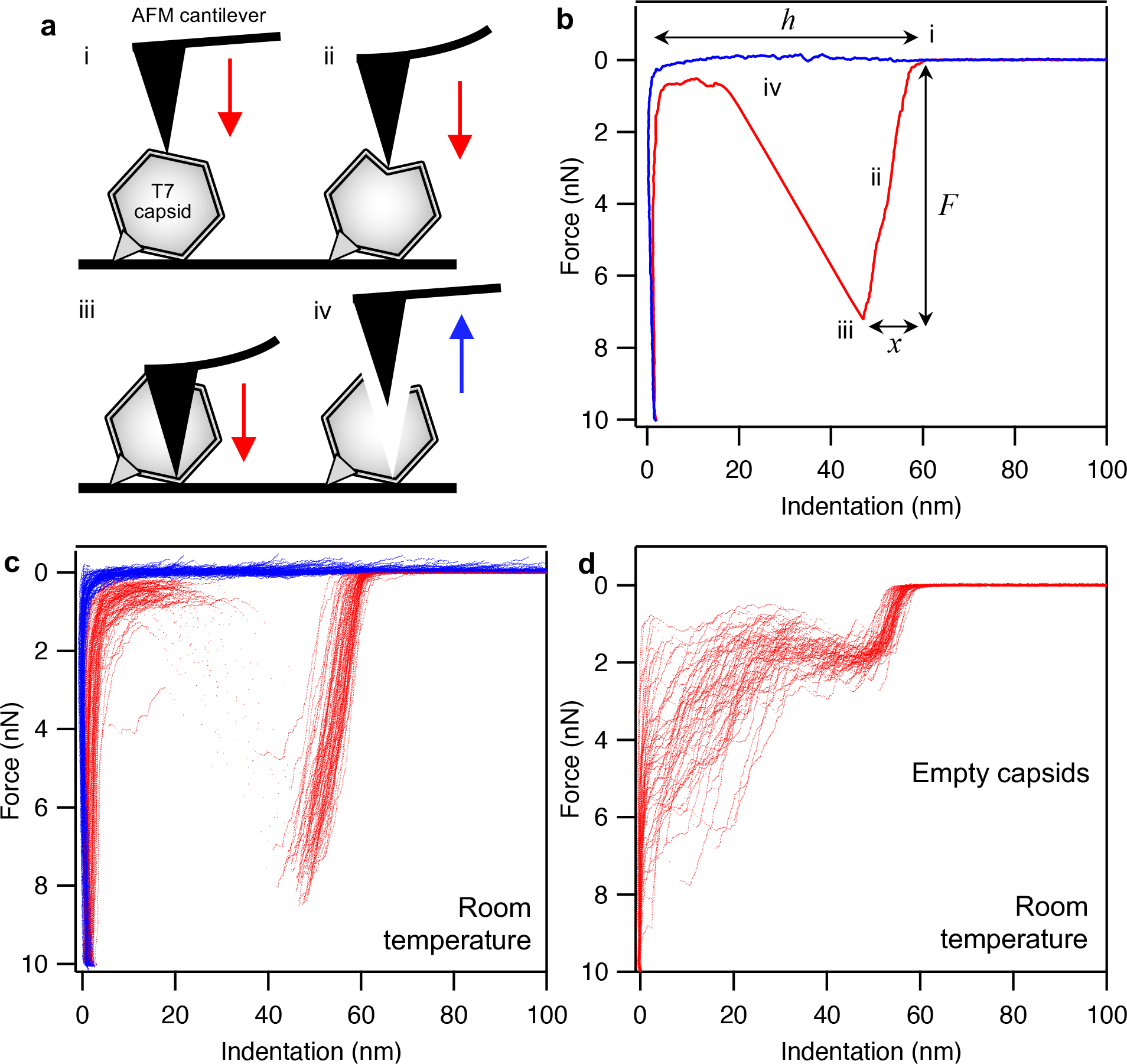
Nanomechanics of T7 phages. **a.** Schematics of mechanical manipulation: the tip of the AFM cantilever is first brought into contact with the T7 phage surface (i) which is then pressed (ii) with a pre-adjusted velocity to 10 nN maximal force during which the capsid eventually ruptures (iii). Finally the cantilever is lifted (iv). AFM cantilever and T7 phage are not to scale. **b.** Representative force *versus* indentation curve obtained at room temperature. Data collected during the indentation half-cycle is displayed in red, whereas those during retraction in blue. Notable stages of the nanomechanics experiments are shown with small Roman numerals (i-iv). Variables extracted from the data (breaking force *F*, maximal indentation distance *x*, capsid height *h*) are shown with italic letters. Capsid stiffness (*k*) is obtained by fitting a line in the initial linear regime of the indentation data (ii). **c.** Dataset containing 80 similar, overlaid force *versus* indentation curves collected in independent experiments on different phage particles at room temperature. Red and blue traces are indentation and retraction half cycles, respectively. **d.** Dataset containing 55 similar, overlaid force *versus* indentation curves (indentation half cycle only), collected at room temperature in independent experiments, which are similar to each other but are distinctively different from the dataset in c (putatively called empty-capsid curves, see later).

**Figure 2.**
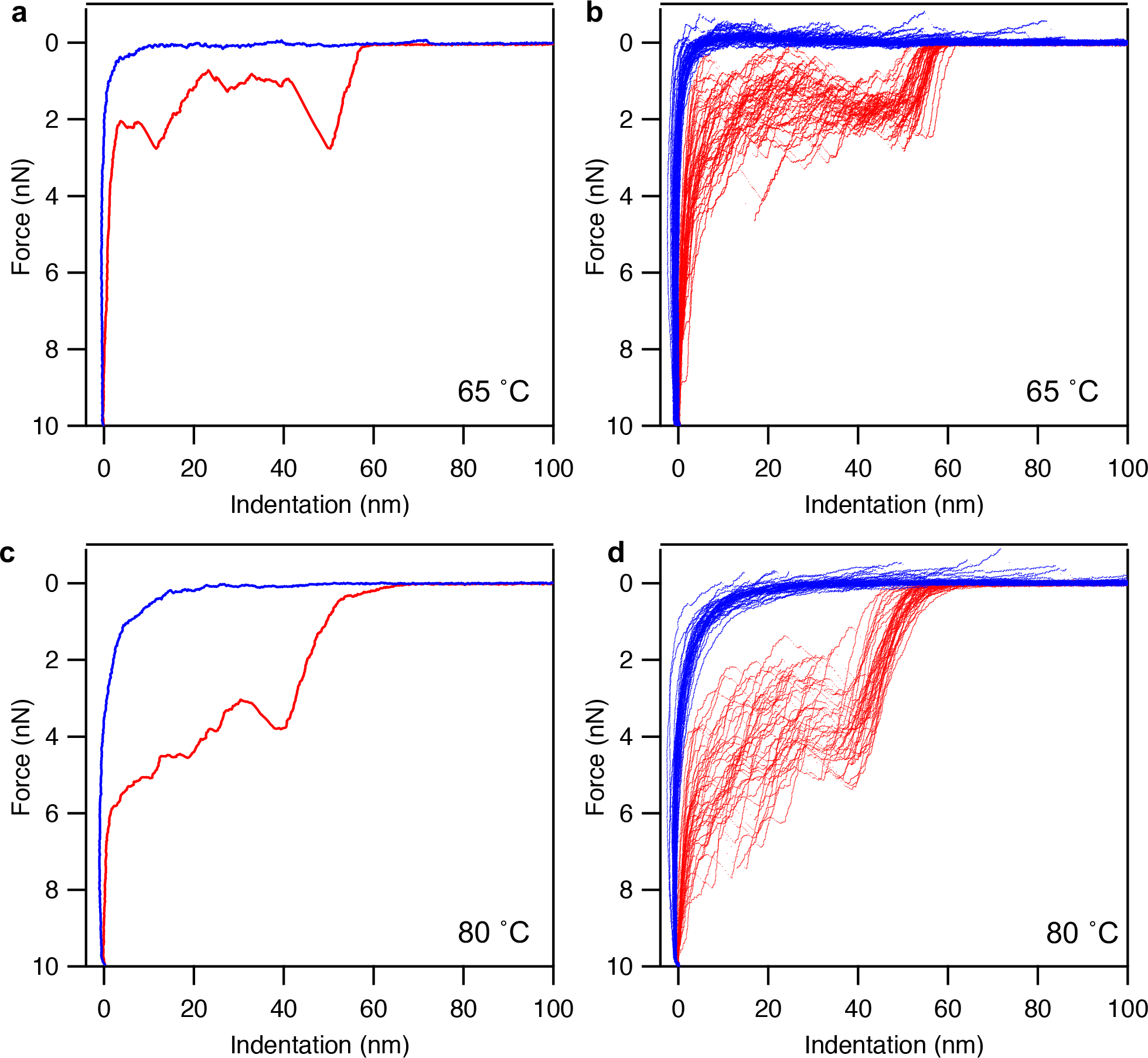
Nanomechanics data of heat-treated T7 phages. Red and blue indicate indentation and retraction half-cycles, respectively. **a.** Representative force *versus* indentation curve measured on a T7 phage particle that has been exposed to a temperature of 65 °C for 15 minutes. **b.** Dataset containing 45 similar, overlaid force *versus* indentation curves collected in independent experiments on different phage particles heat-treated at 65 °C. **c.** Representative force *versus* indentation curve measured on a T7 phage particle that has been exposed to a temperature of 80 °C for 15 minutes. **d.** Dataset containing 41 similar, overlaid force *versus* indentation curves collected in independent experiments on different phage particles heat-treated at 80 °C.

**Table 1.** Nanomechanical and topographical parameters of T7 bacteriophage capsids and globular particles.

|  | RT (DNA-filled) | RT (empty) | 65 °C | 80 °C |
| --- | --- | --- | --- | --- |
| Nanomechanics parameters |  |  |  |  |
| Breaking force ($F$, nN) | $6.90 \pm 0.97$ | $1.90 \pm 0.34$ | $1.61 \pm 0.52$ | $3.83 \pm 1.20$ |
| Capsid stiffness ($k$, $\text{Nm}^{-1}$) | $0.73 \pm 0.12$ | $0.36 \pm 0.10$ | $0.27 \pm 0.11$ | $0.35 \pm 0.15$ |
| Maximal indentation ($x$, nm) | $13.24 \pm 2.13$ | $9.51 \pm 2.41$ | $12.56 \pm 4.70$ | $22.97 \pm 6.61$ |
| Capsid height ($h$, nm) | $60.4 \pm 1.71$ | $59.93 \pm 1.56$ | $59.41 \pm 4.00$ | $60.71 \pm 4.87$ |
| Capsid topographical parameters |  |  |  |  |
| Ratio of tailed capsids | $0.83 \pm 0.1$ | NA | $0.19 \pm 0.12$ | $0.05 \pm 0.06$ |
| Peak capsid height (nm) | $61.8 \pm 3.6$ | NA | $58.2 \pm 1.7$ | $63.3 \pm 5.5$ |
| Capsomer diameter (nm) | $11.0 \pm 0.8$ | NA | $9.8 \pm 1.0$ | $13.9 \pm 2.2$ |
| Globular particle parameters |  |  |  |  |
| Particle height (nm) | $7.6 \pm 3.2$ | NA | $6.9 \pm 2.5$ | $10.0 \pm 5.4$ |

**Fig 3** displays the distribution of the parameters extracted from the force traces (see also **Table 1**). The breaking force values in the RT samples (**Fig. 3.a**) partition into two modes according to the distinct types of force curves (see **Figs. 1.c-d**). The low-force mode of the RT sample aligns well with the histogram peak of the 65 °C data (**Fig. 3.b**). Considering that T7 phages heated to temperatures above 60 °C are thought to loose their DNA (8), we conclude that the low-force peak in the RT data corresponds to empty capsids. The presence of empty capsids in the RT sample indicates that the spontaneous or artificially-induced DNA ejection of T7 phages is not negligible (26). The breaking force is severely reduced in 65 °C-treated capsids (from 6.80 nN to 1.61 nN, see **Table 1**). Since the 65 °C treatment resulted in the release of DNA from the capsids, our results indicate that the presence of packaged DNA within the phage contributes significantly to its mechanical stability. Quite interestingly, the breaking force was increased in the 80 °C-treated T7 phages relative to the ones treated at 65 °C (**Fig. 3.c**). Conceivably, structural rearrangements occurred in the capsomeric proteins between 65–80 °C which resulted in a stabilization of their interactions hence to an increased mechanical stability of the phage particle. Stiffness was largest in the intact T7 phage, and the reduced stiffness values were similar in the RT empty capsids and the heat-treated ones (**Figs. 3.d-f**). Thus, the presence of packaged DNA contributes to the stiffness of the T7 phage. The maximal indentation values progressively increased as a result of heat treatment (**Figs. 3.g-i**), which is a combined effect of the underlying changes in breaking forces and stiffness. Thus, even though the stiffness of 80 °C-treated T7 capsids is reduced, because of the increased breaking forces they withstand greater indentations prior to breakage. The mean capsid height became slightly reduced upon 65 °C treatment (**Fig. 3.j-k**). Notably, the mean capsid height of the empty-capsid RT phages is essentially identical to the 65 °C-treated ones, indicating that the presence of packaged DNA within the phage increases its diameter (by about 10 nm). Conceivably, the DNA pressure inside the phage causes the expansion of the icosahedral phage structure. We note that the 10 nm difference in mean capsid height between the RT and 65 °C-treated capsids is only partly due to the DNA pressure; since capsid height was obtained from mechanical measurements with a pyramidal AFM tip, the presence of upward-oriented phage tails likely shifted the average height to greater values in the RT samples. The capsid height is slightly increased in the 80 °C-treated sample relative to 65 °C, which, as judged from the histogram shape (**Fig. 3.l**), is probably due to the emergence of a subpopulation of capsids with larger diameter.

**Figure 3.**
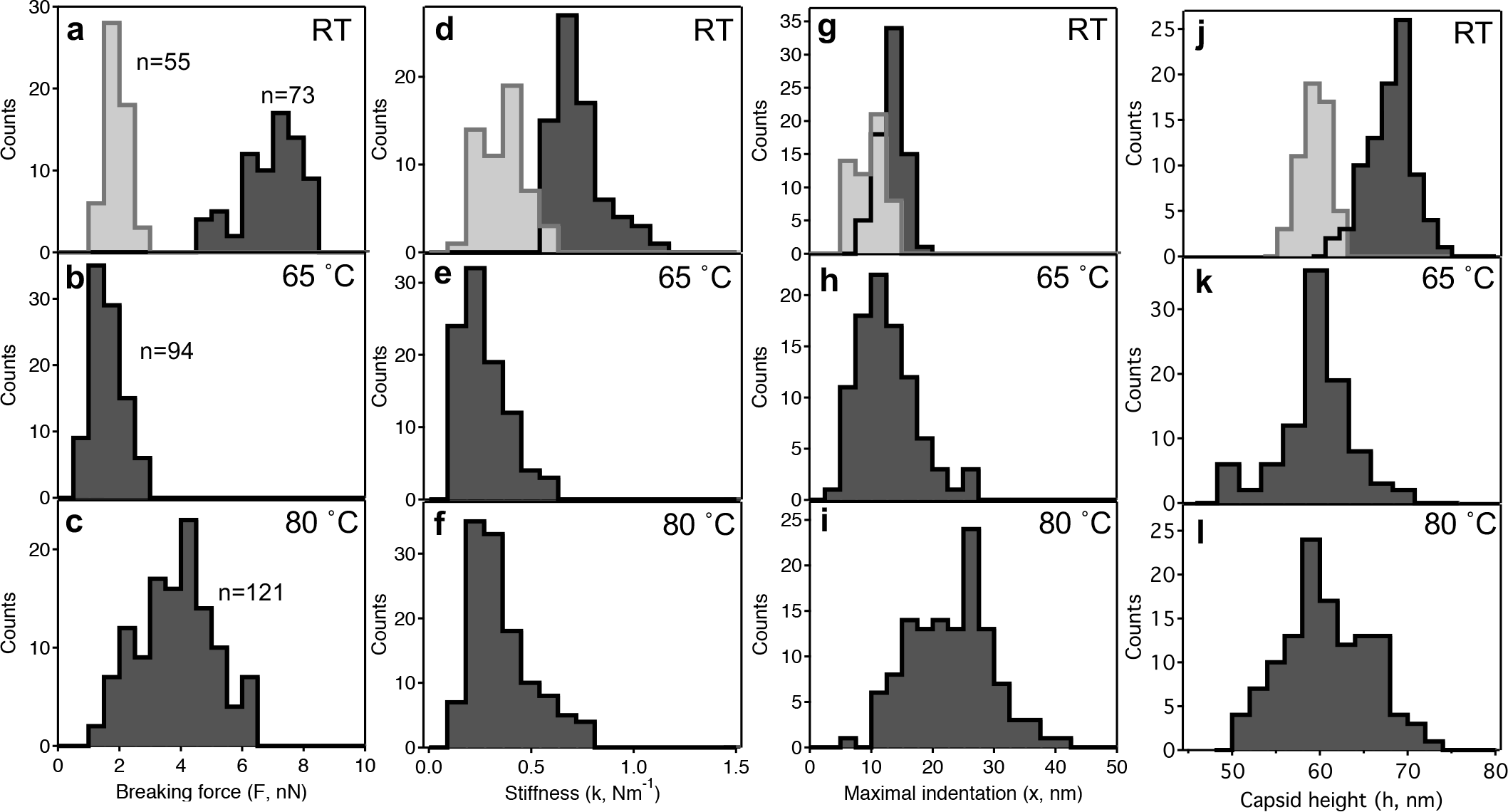
Distribution of variables obtained from nanomechanics data. Breaking force (**a, b, c**), stiffness (**d, e, f**), maximal indentation distance (**g, h, i**) and capsid height (**j, k, l**) histograms for T7 phage particles at room temperature (RT) and ones treated at 65 °C and 80 °C, respectively. Light gray bars correspond to data obtained on empty capsids at room temperature. The numbers (n) refer to the number of force curves analyzed to obtain the nanomechanical parameteres.

### Topographical structure of heat-treated T7 phages

To reveal the structural detail and mechanisms behind the heat-induced nanomechanical changes in T7 capsids, we carried out high-resolution AFM measurements on phage particles exposed to 65 °C and 80 °C (**Fig. 4.a**). In an overview AFM image of a typical RT sample (**Fig. 4.b**) the characteristic T7 phage particles could be visualized against a nearly featureless substrate background. Occasionally, a DNA molecule released from the capsid upon mechanical perturbation could be observed. The mechanically-induced DNA ejection is characterized by the sudden, within-one-scanline appearance of the DNA chain (26). We note that there were a few globular particles in the background, which may correspond to the core T7 phage proteins that become ejected simultaneously with DNA (26). Importantly, the conical tail complex could be observed on most of the phage particles. Depending on the surface binding of the phages the tail complex was oriented in different directions (**Fig. 4.c**). In high-resolution AFM images even the size, the cogwheel shape and the central pore of the capsomeres could be resolved (**Fig. 4.d-e**). Upon 65 °C treatment the topography of the background and to some extent the capsids became different (**Fig. 5**). The most striking feature is that the substrate became covered with a meshwork of DNA chains. A height profile of a section of the background (**Fig. 5.a** **inset**) shows that the cross-sectional height of the individual strands is about 2 nm, which demonstrates that they are indeed DNA. Thus, the 65 °C treatment, as suggested earlier (8), indeed resulted in the release of DNA from the T7 capsids. The second notable feature in the AFM images is that in most of the capsids the conical tail complex is not visible. Even if a tail can be seen, its structure is usually stubby, quite different from a cone (**Fig. 5.c**). Thus, DNA has been released from the phage particles because of a separation of the tail complex from the capsid. Because the gp8 protein plays an important role in the connecting the tail complex to the capsid, we hypothesize that it might be a thermally sensitive component of T7. As a result, large (>10 nm) globular particles can be identified in the background which may correspond to the remnants of the broken-off tail complexes. Although DNA release and the loss of tail were clearly observed in the samples treated at 65 °C, we do not exclude the possibility that these structural transitions may begin to occur at lower temperatures already (27, 28). We note that we were unable to detect the presence of L-shaped tail fibers on the substrate surface. Possibly, the poly-L-lysine-coated surface and the large amount of DNA precluded the binding of the tail fibers in proper orientation. The third striking feature is that the capsid surface became more faceted, and the icosahedron edges and faces emerged more distinctively (**Fig. 5.b**). Such a faceted appearance can be well explained by the shrinkage of the capsids upon DNA release (**Figs. 3.j-k**). In high-resolution AFM images the cogwheel shape of the individual capsomeres could be well identified (**Figs. 5.d, f**). In a few capsids we noticed gaps in the position of the pentameres, which we identify as the exit holes through which DNA escaped (**Fig. 5.e**). The background of T7 samples treated at 80 °C was also densely populated with DNA strands (**Fig. 6.a**). A notable feature in the AFM images is the large number of globular particles scattered in the background. Even large aggregates of the particles could be observed (**Fig. 6.b**). Considering that the size of the aggregates far exceeds that of the tail complex, we hypothesize that the aggregates (hence their component globular particles) originate from the capsid wall. In high-resolution AFM images (**Fig. 6.c-d**) the capsomeres appeared swollen, and they displayed less distinct cogwheel structure according to visual inspection. Altogether, the major transitions of T7 upon heating to 65 °C are DNA release and its associated capsid rearrangements, whereas those upon further heating to 80 °C are structural changes of the capsid wall.

**Figure 4.**
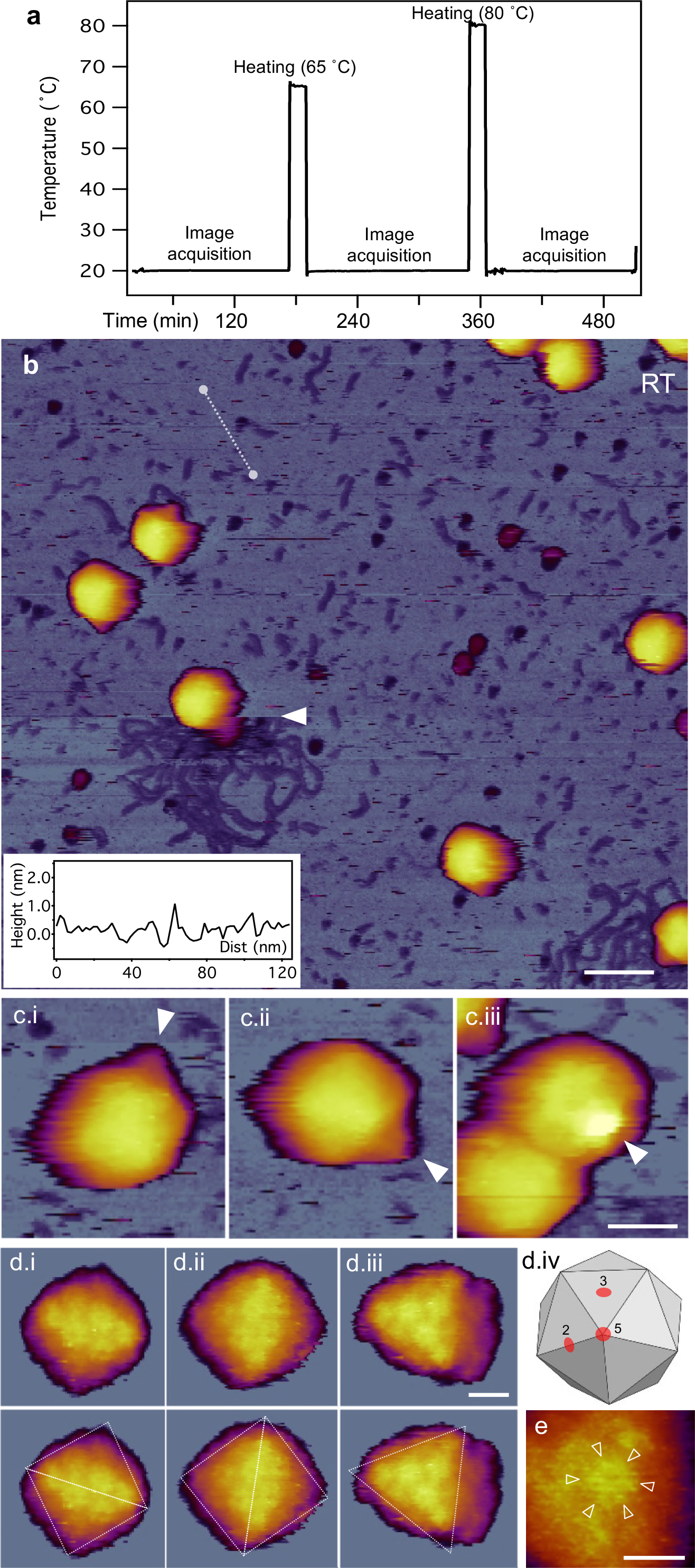
Temperature-dependent AFM measurements on T7 phage particles. **a**. Thermal treatment protocol. Sample temperature *versus* time trace recorded in a typical experiment. The same sample is exposed to consecutive heating (for 15 min) cooling (to 20 °C) and image acquisition (at 20 °C) cycles. **b.** Overview of a 1 μm × 1 μm sample area at room temperature (20 °C). Slow AFM raster scan direction is from top to bottom of the image. White arrowhead points at the nearly instantaneous event of mechanically induced DNA ejection. Scale bar 100 nm. **Inset**, topographical height map along an arbitrarily chosen line in the background (white dashed line). **c.** AFM images of T7 phage particles displaying their conical tail in different orientations. White arrowheads point at the tail apices. Scale bar 30 nm. **d.** High-resolution AFM images of the T7 phage surfaces with resolvable capsomeres. Views along the two-fold (i, ii) and three-fold symmetry axes (iii), which are explained in (iv), are shown. Scale bar 10 nm. In the bottom row dashed guiding lines are superimposed on the respective images to indicate the symmetries. **e.** Magnified view of a cogwheel-shaped hexagonal capsomere. Arroheads point at the spokes of the cogwheel. Scale bar 10 nm.

**Figure 5.**
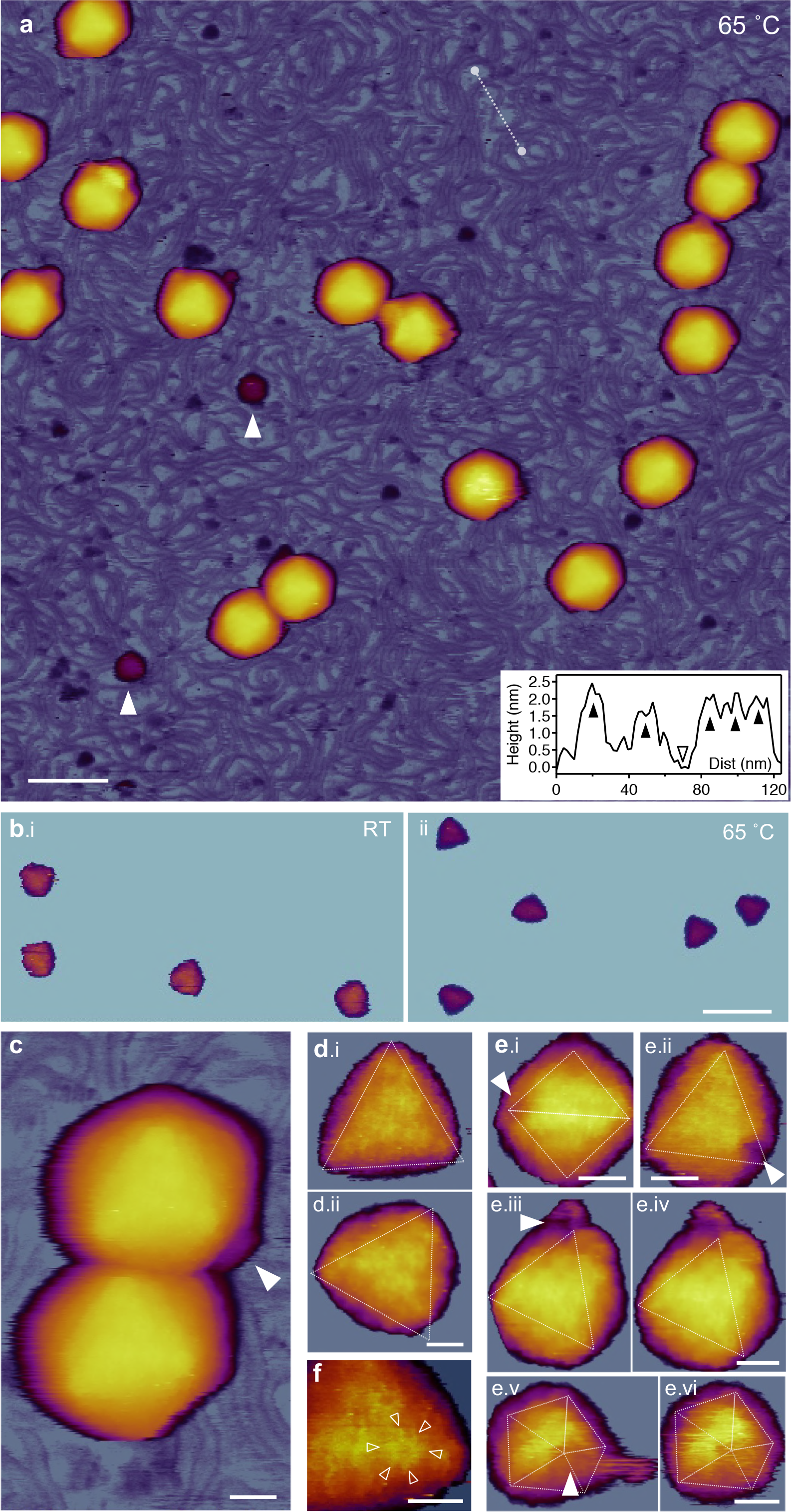
AFM of T7 phages treated at 65 °C. **a.** Overview of a 1 μm × 1 μm sample area. White arrowheads point at large (>10 nm) globular particles. Scale bar 100 nm. **Inset**, topographical height map along an arbitrarily chosen line in the background (white dashed line). Black arrowheads point at DNA cross-sections, whereas the empty arrowhead at the substrate (mica) surface. **b.** Comparison of icosahedral facets of room-temperature (i) and 65-degree (ii) capsids. AFM images contrast enhanced with identical color-scale offset (48 nm) and range (20 nm). **c.** AFM image of two T7 particles. White arrowhead points at the short, stubby tail complex visible on one of the particles whereas there is no visible tail on the other one. Scale bar 20 nm. **d.** High-resolution AFM images of 65 °C-treated T7 phage particles with resolvable capsomeres on their surfaces. Views are along the three-fold symmetry axes. Because of contrast enhancement, only the top facets are visible and the rest of the capsid is hidden. Scale bar 10 nm. **e.** T7 particles with resolvable DNA exit holes (white arrowheads). The exit hole appears as a gap in the location of a missing pentagonal capsomere at one of the icosahedron vertices. Images viewed along the two-fold (i), three-fold (ii, iii, iv) and five-fold symmetry axes (v, vi) are shown. Images iii and v are reconstructed from the rightward fast AFM scanlines, whereas images iv and vi are from leftward (reverse) scanlines from the same sample area. Scale bars, 20 nm. **f.** Magnified view of a cogwheel-shaped hexagonal capsomere. Arroheads point at the spokes of the cogwheel. Scale bar 10 nm.

**Figure 6.**
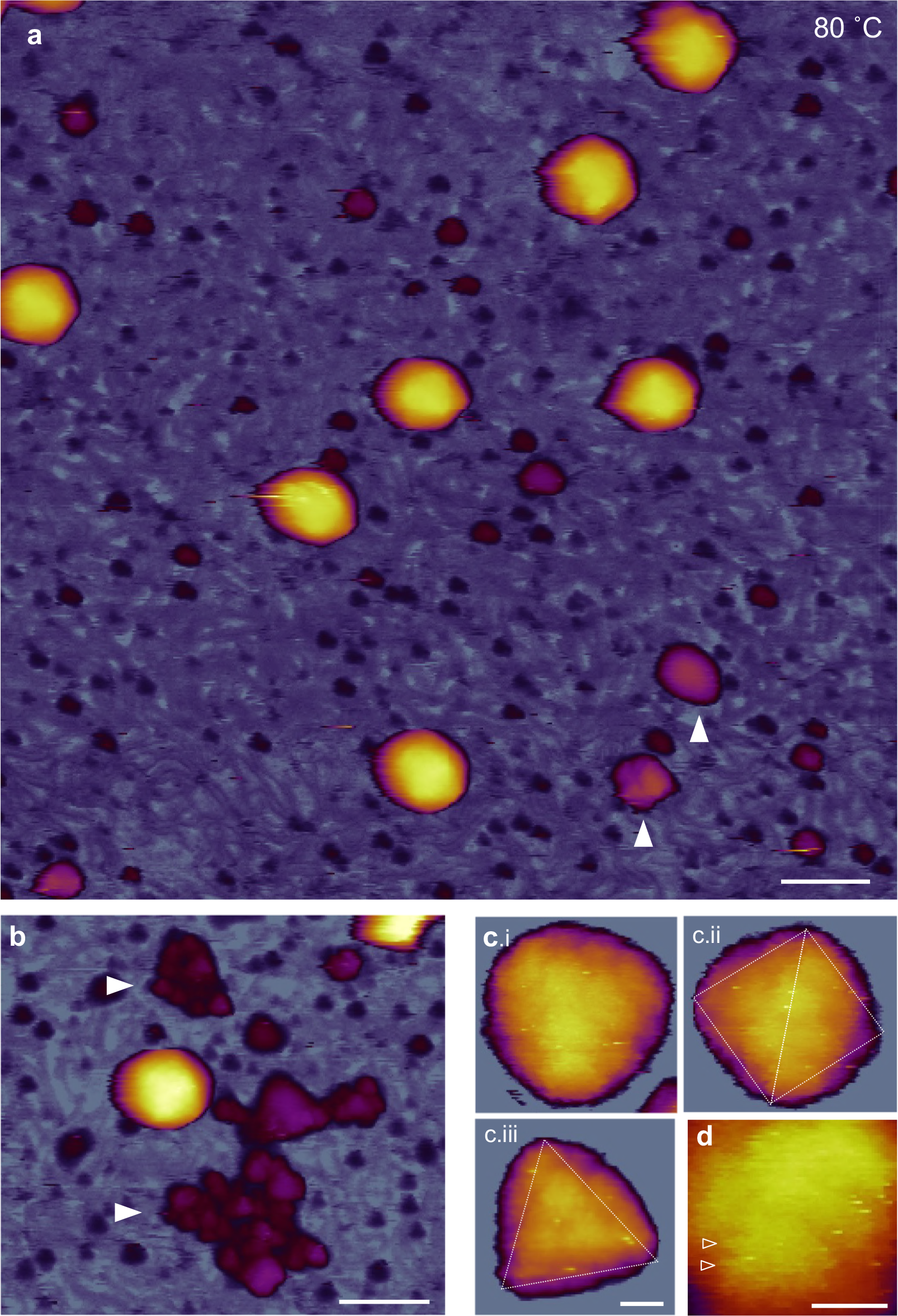
AFM of T7 phages treated at 80 °C. **a.** Overview of a 1 μm × 1 μm sample area. White arrowheads point at large (>10 nm) globular particles. Scale bar 100 nm. **b.** AFM image showing large aggregates of globular particles (white arrowheads) **c.** High-resolution AFM images of 80-degree-treated T7 phage particles with resolvable capsomeres on their surface. Views along the three-fold (i, iii) and two-fold symmetry axes (ii) are shown. Scale bar 10 nm. **d.** Magnified view of the capsomeric structure. Arroheads point at putative spokes of the originally cogwheel-shaped capsomere. Note that the central pore cannot be resolved, most likely due to the swelling of the protein matrix. Scale bar 10 nm.

**Figs 7**-**8** show the analysis of topographical data. Representative height profiles across individual capsids (**Fig. 7.a**) demonstrate the heat-induced topographical changes in T7. Upon 65 °C treatment the capsid slightly shrunk, its faces became flattened and its edges more distinct. The 80 °C-treated capsid shown here became swollen and its surface rugged. The ratio of capsids with visible tail complexes progressively reduced with heat treatment (**Fig. 7.b**). The capsomere diameter considerably increased upon 80 °C treatment (**Fig. 7.c**). We hypothesize that thermally-induced conformational changes, most likely partial denaturation, has occurred in the gp10A capsomeric proteins which resulted in an increase in their apparent volume. It might well be possible that the partial denaturation exposed the hydrophobic core of the capsomere proteins, and a hydrophobic interaction occurred in between neighboring capsomeres. Such an interaction may explain the increase in breaking force between 65 °C and 80 °C observed in the nanomechanical experiments (**Figs. 3.b-c**). The peak capsid height decreased slightly upon 65 °C treatment (**Figs. 7.d-e**), but the 80 °C treatment resulted in the emergence of a sub-population with larger height values (**Fig. 7.f**). We hypothesize that this sub-population corresponds to capsids with swollen wall structure. The peak capsid height analysis shown here more-or-less reflects the tendencies observed in the nanomechanical experiments. However, because AFM images allow us to select the tallest topographical point on the capsids, the peak height analysis is more sensitive to local variations which are hidden or averaged out in the nanomechanics experiment. The number of globular particles progressively increased in the samples upon heat treatment (**Fig. 8.a**). The height of the major population of the particles is centered around 6 nm regardless of heat treatment (**Figs. 8.b-d**). In the 80 °C-treated samples particle populations with much larger heights emerged (**Fig. 8.c**). While the ~6 nm particles may correspond to the ejected core proteins, the large globular particles are most likely capsomeric proteins and their aggregates which appear due to the complete disassembly of some of the capsids.

**Figure 7.**
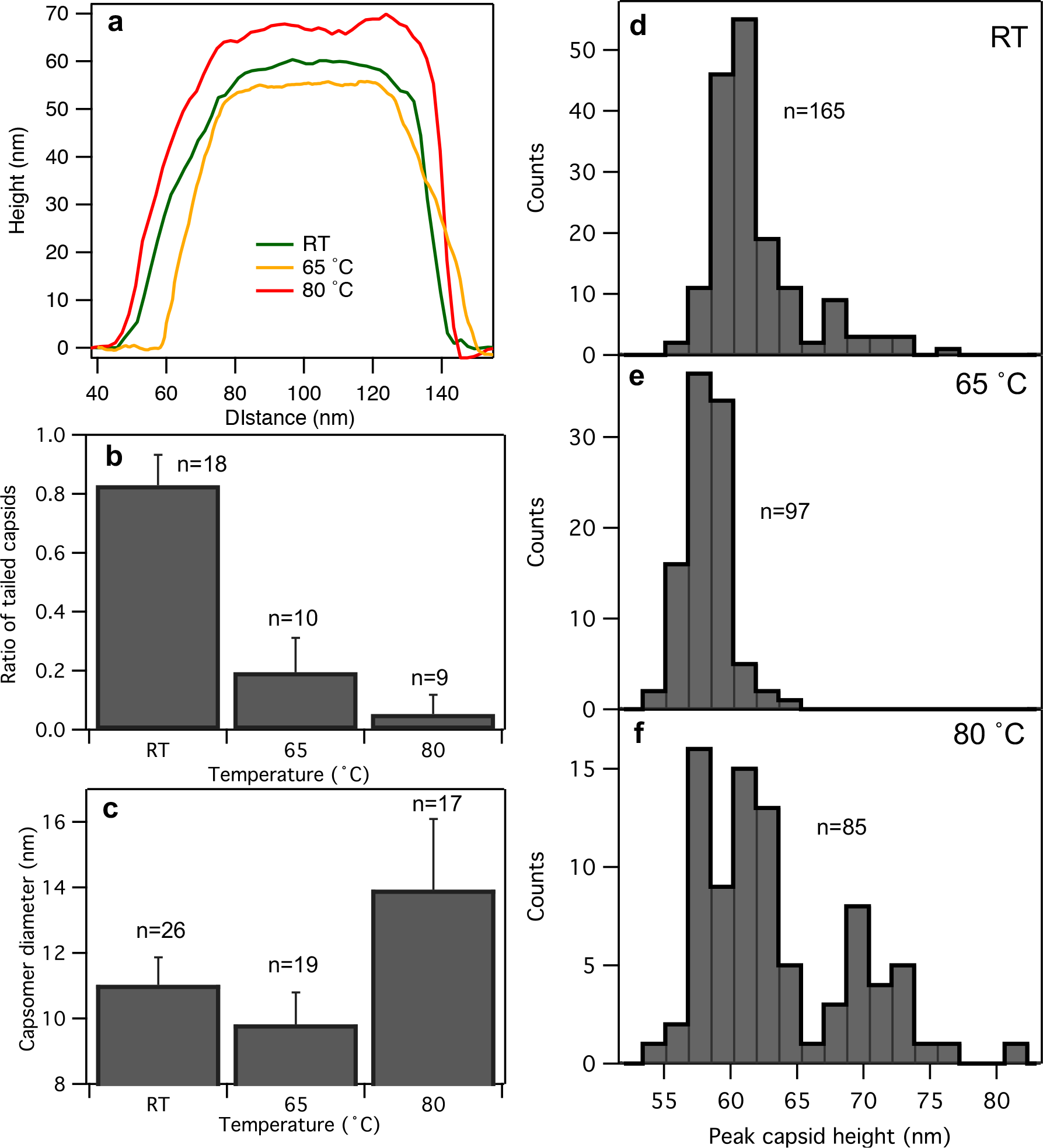
Analysis of capsid topography data. **a.** Topographical height map along the cross-section of either a capsid at room temperature (green trace) or ones treated at 65 °C (orange) or 80 °C (red). **b.** Ratio of capsids with tails as a function of temperature. The numbers above the bars represent the number of fields analyzed for every T7 particle. Error bars represent standard deviation (SD). **c.** Capsomer diameter as a function of temperature. The numbers above the bars represent the number of capsomers measured. Error bars represent standard deviation (SD). **d, e** and **f** show histograms of peak capsid heights for room-temperature (RT), 65 °C and 80 °C treated T7 phages, respectively. Peak height refers to the tallest topographical point in the capsid image. The numbers (n) refer to the number of T7 phage particles analyzed.

**Figure 8.**
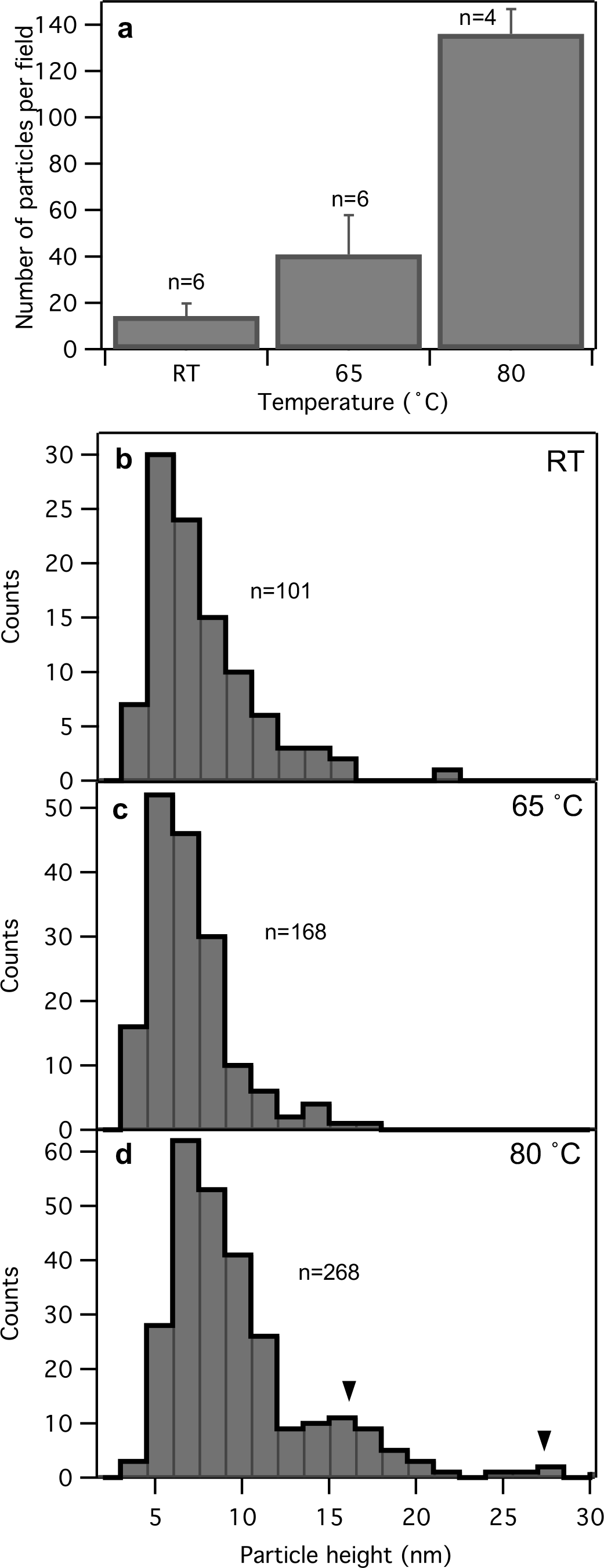
Analysis of topographical data of globular particles. **a.** Number of globular particles per field as a function of temperature. The numbers above the bars represent the number of fields analyzed for every particle. Error bars represent standard deviation (SD). **b, c** and **d** show histograms of globular particle height for samples at room temperature (RT) and ones treated at 65 °C and 80 °C, respectively. Black arrowheads point at populations of large globular particles. The numbers (n) refer to the number of globular particles analyzed.

### Model of thermally-induced structural changes in T7

We propose the following phenomenological model to explain our observations (**Fig. 9**). At room temperature (**Fig. 9.a**) T7 displays a characteristic icosahedral structure with distinctive conical tail complex. The icosahedron is slightly swollen due to the DNA pressure inside the capsid. The DNA-filled T7 phages have a high stiffness and withstand (instantaneous) forces up to about 8 nN prior to breakage. Upon heating to 65 °C (**Fig. 9.b**), the genomic DNA is ejected from the capsid. The release of DNA is caused by the conical tail complex breaking off the capsid, rather than by activating the natural DNA-ejection machinery. It is currently unclear whether the entire genome of T7 exits the capsid or a portion remains inside. It is hypothesized that during its natural DNA ejection, only part of the T7 genome is driven out of the capsid due to the DNA pressure. The remaining DNA is thought to be pulled into the *E.coli* by an active process (29–31). However, because the thermally-induced changes involve the loss of the entire tail complex, there might be enough room for the nearly complete release of the T7 genome. Regardless of how much DNA exits during this process, the resulting drop in DNA pressure leads to a shrinkage and a more faceted appearance of the capsid. Upon further heating to 80 °C (**Fig. 9.c**) the capsids do not disappear, but they are still present with a maintained global structure. A partial denaturation likely takes place in the gp10A proteins that form the capsomeres and hence the capsid wall. The partial protein denaturation within the global confinement of the capsid architecture and the resulting exposure of hydrophobic protein regions result in capsomere swelling and a new set of inter-capsomeric interactions, most probably *via* hydrophobic protein regions. Facilitated folding and misfolding following repetitive partial or complete denaturation have been observed in proteins confined either in a chaperonin system (32) or in a force field (33). Notably, capturing a protein in the misfolded state results in a considerable conformational expansion (34). We envision that similar processes may occur in the gp10A proteins heated to 80°C then cooled to room temperature. In the end, the capsid wall becomes thicker, and the entire capsid surface becomes rugged. It is quite conceivable, that the hydrophobic stabilization is not the result of the heating *per se*, but of the relaxation from the thermal exposure. That is, capsids that did not completely fall apart during the 80 °C treatment may relax into a stabilized structure upon cooling back to room temperature.

**Figure 9.**
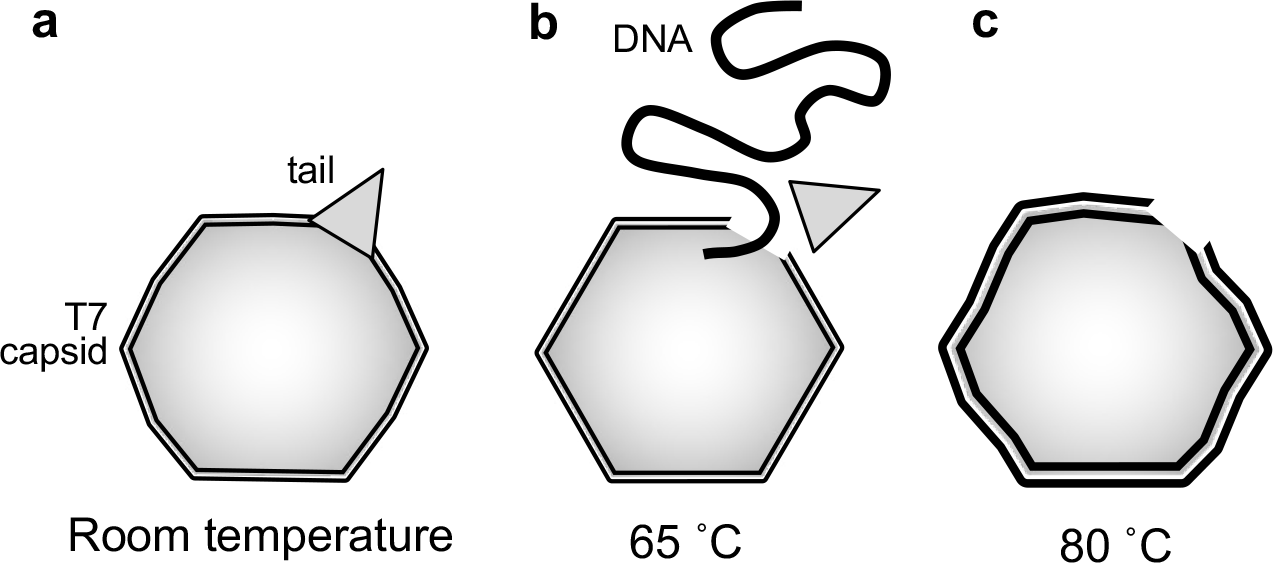
Schematic model of thermally-induced changes in the T7 bacteriophage. At room temperature (**a**) the capsid is slightly swollen because of the DNA pressure inside. The bulging of the capsid wall as shown in the scheme is not to scale. Upon heating to and incubating at a temperature of 65 °C (**b**), the tail complex is broken off, resulting in the release of the genomic DNA. The capsid becomes more faceted due to the relaxation of the capsid pressure. Finally, at 80 °C (**c**) the capsid becomes swollen and its surface irregular, and the capsids may become fragmented into large globular particles (this last step is not shown).

### Conclusions

We have directly shown that exposing T7 to a thermal treatment at 65 °C caused the release of its genomic DNA due to the tail complex breaking off the capsid. The loss of DNA results in a relaxation of the capsid structure accompanied by a reduced stiffness and breaking force. Further heating to 80 °C leads to rearrangements within the capsid wall caused most likely by partial denaturation of the component gp10A proteins. Even though the loss of DNA destabilizes the capsids, they are remarkably able to withstand high temperatures with a more-or-less intact global topographical structure. Thus, partial denaturation within the global structural constraints of the viral capsid may have a stabilizing effect. Therefore, understanding the structural design of viruses may help in constructing artificial nanocapsules for the packaging and delivery of materials under harsh environmental conditions.

## Materials and methods

### T7 preparation

T7 (ATCC 11303-B7) was grown in *Escherichia coli* (ATCC 11303) host cells and purified according established methods (35). Briefly, the phage suspension was concentrated on a CsCl gradient and dialyzed against buffer (20 mM Tris-HCl, 50 mM NaCl, pH 7.4) (10). T7 bacteriophage concentration was determined from optical density by using an exctinction coefficient of ε_260_ = 7.3 × 10^3^ (mol nucleotide bases × L^−1^ × cm^−1^). The dialyzed T7 samples were kept at 4 °C for a few months without significant loss of activity. Prior to further use the T7 samples were diluted with PBS (137 mM NaCl, 2.7 mM KCl, 10 mM Na_2_HPO_4_, 1.8 mM KH_2_PO_4_, pH 7.4).

### Atomic force microscopy and nanomanipulation

T7 samples properly diluted in PBS were applied to freshly cleaved mica functionalized with glutaraldehyde (25, 36). The dilution was adjusted so that an approximate surface density of 10 phage particles per μm^2^ is achieved. A freshly-cleaved mica was first incubated with poly-L-lysine (0.01 % aqueous solution) for 20 minutes at room temperature, then rinsed extensively with MilliQ water and dried with a stream of high-purity N_2_ gas. Subsequently, the surface was incubated with 10 % aqueous glutaraldehyde for 30 minutes at room temperature, then rinsed extensively with MilliQ water and dried with a stream of high-purity N_2_ gas. Finally, a sample of T7 phage was loaded onto the substrate surface and incubated for 40 minutes on ice. Unbound viruses were removed by gentle washing with PBS. Non-contact mode AFM images were acquired with an Asylum Research Cypher instrument (Asylum Research, Santa Barbara, CA) by using silicon-nitride cantilevers (Olympus BL-AC40TS-C2 or Nanoworld PNP-TR). 512 × 512-pixel images were collected at a typical scanning frequency of 0.3–1.5 Hz and with a mean indentation force of about 30 pN. All of the images presented in this work were collected on non-fixed samples under aqueous buffer conditions. For temperature-dependent measurements we used the cooler/heater stage of the AFM instrument. Temperature was kept constant with a precision of 0.1 °C. Evaporation of water was prevented by the sealed container housing the AFM scanner. For nanomechanical measurements the surface-bound viruses were manipulated by first pressing the cantilever (Nanoworld PNP-TR, lever 1) tip against the apex of the virus, then pulling the cantilever away with a constant, pre-adjusted rate (25). Typical cantilever movement rate was 1 μm/s except where noted otherwise. Stiffness was determined for each cantilever by using thermal method (37).

### Image processing and data analysis

AFM images and force spectra were analyzed using algorithms built in the Cypher controller software (AsylumResearch, Santa Barbara, CA). Indentation distance (*z*) was calculated from cantilever displacement (*s*), force (*F*) and cantilever stiffness (*k*) as

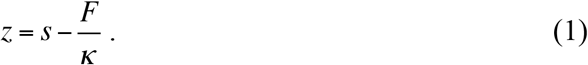

AFM images were corrected for flatness of field (within a few Å) and their color contrast was adjusted in order to better communicate the relevant features. No additional image processing was carried out.

### Statistics

The number of nanomechanical curves, images and particles (T7 or globular) analyzed are shown in the relevant figures. The results shown in this manuscript were collected in 15 independent nanomechanics and 9 independent AFM imaging experiments. Statistical analyses and graph plotting was carried out by using either KaleidaGraph (v.4.5.1, Synergy Software, Reading, PA) or IgorPro (v. 6.34A, Wavementrics, Lake Oswego, OR) programs.

## Author contributions

Z.V. performed research, analyzed data, wrote paper; G.C. contributed analytic tools; L.H. contributed analytic tools; M.K. designed research, performed research, analyzed data, wrote paper.

## Acknowledgements

This work was supported by grants from the Hungarian National Research, Development and Innovation Office (K109480; K124966; VKSZ_14-1-2015-0052; NVKP-16-1-2016-0017 National Heart Program). The research leading to these results has received funding from the European Union’s Seventh Framework Program (FP7/2007-2013) under grant agreement n° HEALTH-F2-2011-278850 (INMiND).

